# Fusion peptides of enveloped viruses actively mediate membrane fusion in model cell membranes

**DOI:** 10.64898/2026.06.05.729768

**Authors:** Mahsa Mohammadian, Ralf Seemann

## Abstract

Enveloped viruses can enter host cells by fusing their membrane with that of the host cell, a process known as membrane fusion. This process depends on specific fusion proteins located on the viral particle surface, which contain a short, relatively hydrophobic segment called “fusion peptide” that binds to the host membrane. To investigate the fusion efficiency of various fusion peptides, we create simplified non-infectious virus like particles decorated with different fusion peptides and fuse them with an artificial cell membrane. For this purpose, microfluidic devices are used to create supported lipid bilayers while the result of the fusion process is studied by fluorescence microscopy. Our study provides structural insights into the interactions between virus particles and cell membranes, which can facilitate the development of new therapeutic strategies and more effective viral vectors for therapeutic applications.

## Introduction

Viral infection proceeds by binding of the virus to the host cell membrane, followed by cell entry, allowing the virus to deliver its genetic material to initiate genomic replication (*1,2*). Some viruses, particularly some enveloped viruses, enter the cells via two possible pathways: direct fusion with the plasma membrane or entry via endocytosis and subsequent fusion within the endosome (*3, 4*). Although in both pathways, membrane fusion is the critical step that allows the viral genome to cross a cellular membrane barrier, the latter process typically requires a low pH environment to trigger the fusion mechanism (*5*).

During fusion, two distinct lipid bilayers combine into a single continuous membrane, allowing their internal contents to mix, a process that is also essential for many other biological functions (*6*). To mediate the fusion process, enveloped viruses such as influenza, HIV, coronaviruses, etc. use specialized surface proteins known as fusion proteins, which facilitate fusion between their viral envelope and the host cell membrane (*1, 4*). Fusion proteins are typically single-pass integral membrane proteins with relatively large ectodomains (*1, 7*).

A key component of the fusion proteins is the fusion peptide, a short -usually 20 to 30 residues in length-, relatively hydrophobic sequence typically located at or near the N-terminus of the viral fusion protein (*1, 8, 9*). In the resting (non-fusogenic) state of the fusion protein, the fusion peptide is buried in a hydrophobic pocket of the soluble ectodomain. Upon activation, such as membrane apposition or a drop in pH, the protein undergoes a conformational change that exposes the fusion peptide, allowing it to insert into the host membrane (*10*). Fusion peptides are often enriched in glycine and alanine residues (*1, 11*), a composition that provides considerable conformational flexibility and structural polymorphism. This enables them to adapt to different lipid compositions, pH conditions, and protein environments. For instance, the HIV fusion peptide can adopt either an *α*-helical or *β*-sheet conformation depending on the cholesterol content of the membrane (*1*).

The essential role of fusion peptides in membrane fusion has been demonstrated through numerous mutagenesis studies, where deletion or even minor single amino acid substitutions can significantly disrupt the protein’s ability to mediate fusion (*12*). Interestingly, mutations in other regions of the fusion protein generally tend to have a less severe impact on its function, underscoring the critical role of the fusion peptide (*13, 14*).

While the primary role of fusion peptides in membrane binding has been well established, recent research suggests that their function extends beyond simple membrane insertion. Emerging evidence indicates that fusion peptides may actively participate in the restructuring and destabilization of lipid bilayers, influencing the kinetics and efficiency of the fusion process (*15*). When fusion peptides insert into a target bilayer they induce local membrane deformations that facilitate the merging of the viral and host membranes (*1,8*). These findings challenge the traditional view that fusion peptides serve merely as passive anchors, highlighting their potential as active modulators of membrane fusion. Infections can be avoided by preventing the fusion of the viral envelope with the cell or endosomal membrane. Likewise, targeting the viral fusion protein with a vaccine can serve as an effective preventive measure. For instance, influenza vaccines work by targeting the virus fusion proteins (*3*).

In this work, we investigate the lesser-known functions of fusion peptides, particularly their active involvement in the fusion process between viral and host membranes. To achieve this, simplified model membranes and microfluidic platforms were employed to study fusion processes under controlled experimental conditions. Analyzing the experimental data demonstrates that fusion peptides do more than merely binding to the host membrane, they actively mediate fusion. These insights could contribute to the development of novel therapeutic strategies and more efficient viral vectors for medical applications.

## Results and Discussion

### Lipid Bilayer Characterization

Supported lipid bilayers (SLBs) were formed within a microfluidic device using Cy5-labelled large unilamellar vesicles (LUVs) with a lipid composition inspired by the major lipid classes and cholesterol content of mammalian plasma membranes (Table 1), consisting of DOPC, DOPE, DOPS, sphingomyelin (SM), cholesterol (Chol) at a molar ratio of 30:18:6:6:40 (mol%) (*16,17*). Bilayer formation was monitored in real time by fluorescence microscopy. During the initial stages, heterogeneous and punctate fluorescence patterns were observed, corresponding to partially fused vesicles on the substrate (Fig. 1a), indicative of an intermediate assembly state. With continued incubation and removal of excess vesicles, the fluorescence signal became homogeneous across the surface, consistent with the formation of a continuous and defect-free SLB (Fig. 1b). A schematic representation of the resulting supported bilayer is shown in Fig. 1c.

**Table 1:** Lipid composition of LUVs that were used to (a) form a supported lipid bilayer (mimicking Plasma Membrane) and (b) fabricate VLPs (mimicking Viral Membrane).

| Composition of the SLB |  | Lipidic composition of the VLP |  |
| --- | --- | --- | --- |
| Lipid | Molar ratio | Lipid | Molar ratio % |
| DOPC | 30 | DOPE | 48 |
| DOPE | 18 | DOPC | 9 |
| DOPS | 6 | DOPS | 25 |
| SM | 6 | DSPA | 3 |
| Chol | 40 | SM | 15 |
| Cy5-PC | 2 | Rhod-PE | 2 |

**Figure 1:**
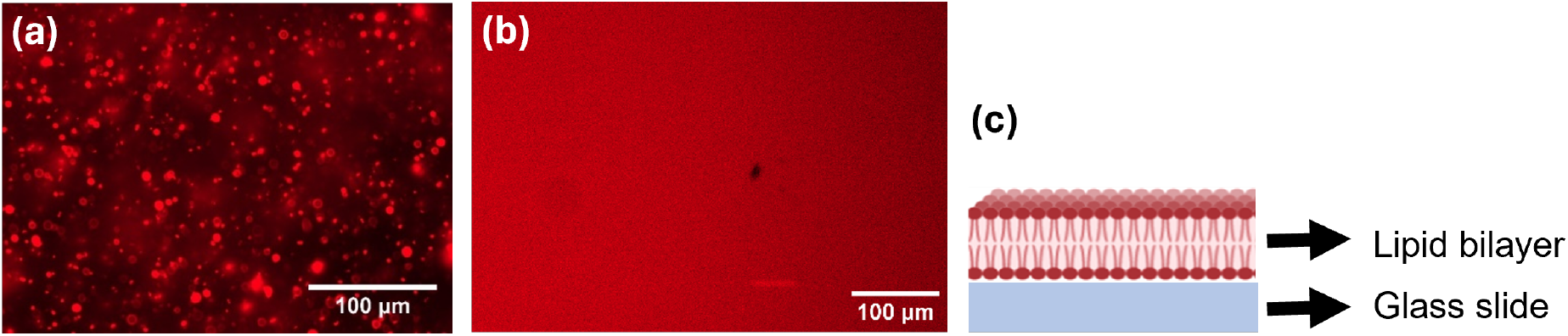
Formation and visualization of a supported lipid bilayer with the composition of DOPC:DOPE:DOPS:SM:Chol (30:18:6:6:40 Mol%). (a) Fluorescent micrograph of an intermediate step during bilayer formation and (b) final stage of a formed lipid bilayer in top-down view, labeled with Cy5-PC and observed with a 20x objective, (c) schematic image of the supported bilayer on a glass slide, side view.

Fluorescence recovery after photobleaching (FRAP) experiments were performed to verify bilayer integrity and quantify lateral lipid mobility of the finally formed SLBs. Representative FRAP images show the bleached region and its subsequent fluorescence recovery over time (Fig. 2a). The corresponding recovery curve is shown in Fig. 2b and is analyzed using the Soumpasis model (*18, 19*). Fitting of the recovery data yielded a lipid diffusion coefficient of approximately 0.47 µm^2^*/*s. This value indicates moderate lateral mobility within the bilayer and is consistent with previously reported diffusion coefficients for cholesterol-containing supported lipid systems. *Bacia et al*. reported diffusion coefficients of (0.3–0.5) µm^2^*/*s for cholesterol-rich membrane phases (*20*), while *Zhang et al*. observed a decrease in lipid mobility from 1.3 µm^2^*/*s in pure DOPC bilayers to 0.45 µm^2^*/*s upon incorporation of 30 mol% cholesterol (*21*). Overall, our measurements confirm the successful formation of a continuous, fluid, and mechanically stable SLB. The obtained diffusion characteristics are in agreement with expectations from the literature and further indicate that the membrane system is physiologically relevant and provides a robust and reproducible platform for investigating virus-membrane interactions.

**Figure 2:**
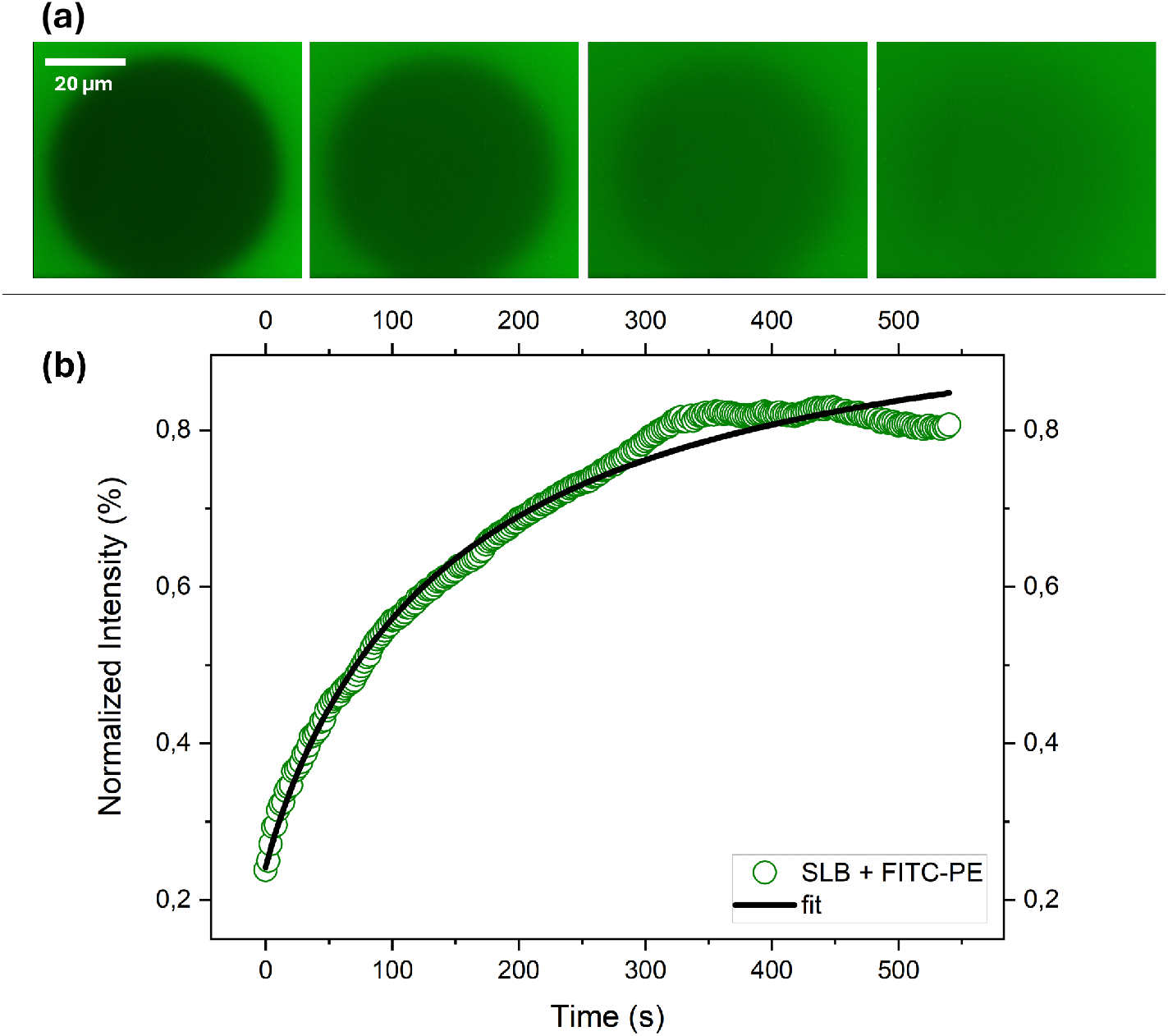
FRAP characterization for FITC-labeled SLB composed of DOPC:DOPE:DOPS:SM:Chol (30:18:6:6:40 Mol%) (a) Representation of the FRAP experiment on SLB. First image on the left shows the bleached area of the SLB and the other images shows the recovery steps of the same spot. (b) Fluorescence recovery curve from FRAP experiments together with the Soumpasis model fit.

### Virus Like Particle-Bilayer Fusion Experiment

To study the membrane fusion between viruses and host cell membranes, we combined a microfluidic membrane fusion assay with model lipid bilayers mimicking viral and host membrane compositions to quantitatively assess how different fusion peptides modulate fusion efficiency (*16, 17, 22*). Fusion peptides from different enveloped virus families, and a human sourced peptide that is known as non-binder (as control sample) were chosen. Membrane composition of virus like particles (VLPs) and the peptide sequences used throughout this work are summarized in Table 1–2. Fusion efficiency of virus like particles (VLPs) was quantified by monitoring the increase in Rhodamine fluorescence intensity upon VLP contacts with the bilayer. Analysis was performed in a defined region of the microfluidic channel (100 µm *×* 100 µm), which was initially homogeneously covered by a supported lipid bilayer, confirmed by uniform Cy5 fluorescence. Fluorescence intensities were extracted from the microscopy images using Zeiss Zen 3.3 (blue edition) and further processed in Origin.

**Table 2:**
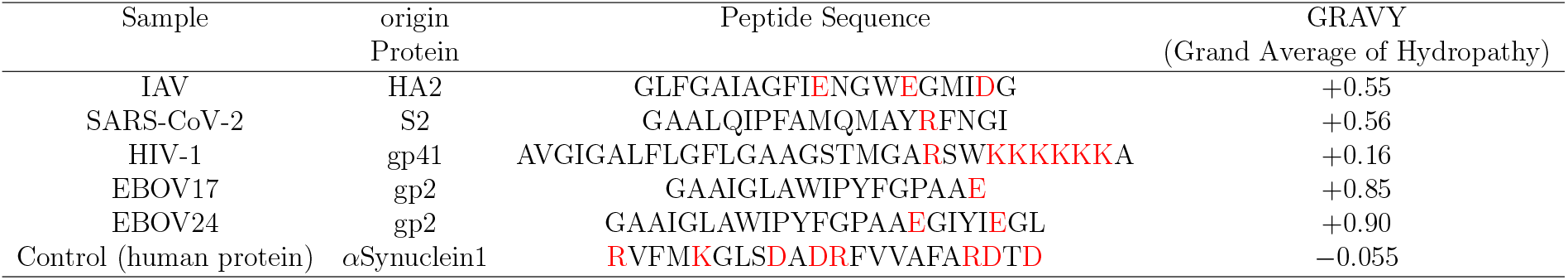
Respresentation of the studied enveloped viruses, their origin protein, corresponding fusion peptide sequences and their hydropathy scale, GRAVY (Grand Average of Hydropathicity) calculated using Kyte-Doolittle method. Charged residues are highlighted in red.

Representative fluorescence images across increasing contact times with VLP dispersions (flushing cycles) are shown in Fig. 3a. The temporal evolution of the averaged fluorescence intensity for different virus fusion peptides (IAV, EBOV17,EBOV24, HIV-1 and SARS-CoV-2), a non-binder (*α*Synuclein1, *α*Syn1) and bare lipid bilayers as control tests, were obtained and presented in Fig. 3b. Already by visual inspection it is obvious that the fusion activity of the various viral peptides is higher than that of the two control tests, whereas difference between the various viral peptides are rather small and become clear only after longer contact times. To enable a quantitative comparison of the fusion efficiencies, two complementary analytical approaches were applied. In the first approach, the increasing fluorescence intensity was described fitting an exponential model (*y* = *B ·* (1 − exp(−*A · t*))) to each individual data set, from which the initial slope at *t* = 0 was derived and averaged across repeated experimental realizations. In the second approach, the early interaction phase was independently evaluated using a linear fit (*y* = *a · t* + *b*) applied to the initial three contact time intervals of each data set, and the resulting slopes were likewise averaged over five experiments. Figure 3c presents an example of both applied fitting models for an individual IAV fusion peptide dataset. A comparative summary of the averaged fusion efficiencies derived from both approaches is provided in Fig. 4.

**Figure 3:**
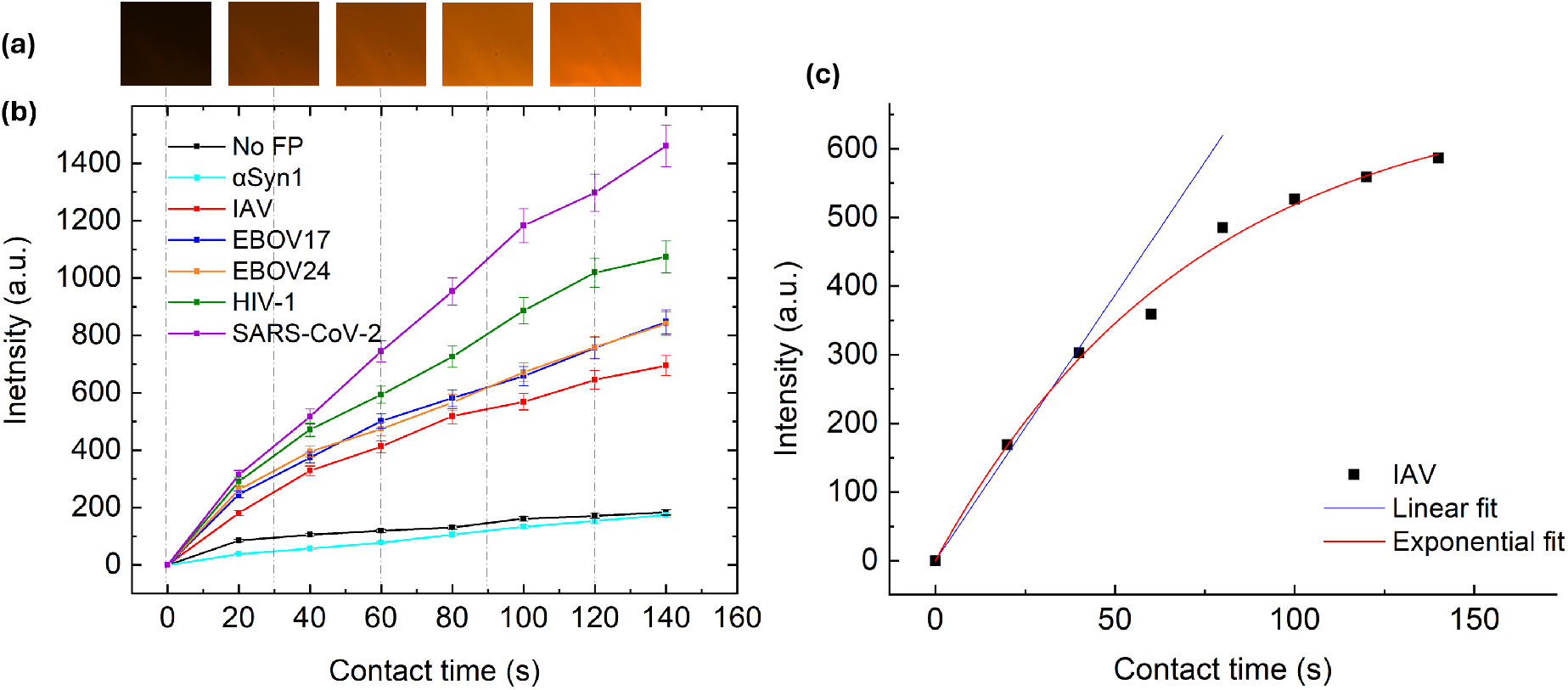
Time-dependent fluorescence intensity of the supported lipid bilayer, following interaction with VLPs. (a) Representative fluorescence microscopy images showing increasing fluorescent signal of the SLB after fusion with Rhod-labelled VLPs. (b) Mean fluorescence intensity from fluorescent images such as shown in panel (a), averaged over five independent experiments, plotted against the VLP-bilayer contact time. Different colours and symbols represent individual VLP or control conditions. Error bars correspond to the standard deviation obtained from five independent measurements. (c) Example analysis of fluorescence intensity data for IAV fusion peptide with the supported lipid bilayer. Exponential (red; *y* = *B* (1 − exp(−*A · t*))) and linear (blue; *y* = *a · t* + *b*) fits were applied to the results of a single experiment (*R*^2^ ≈ 0.99).

**Figure 4:**
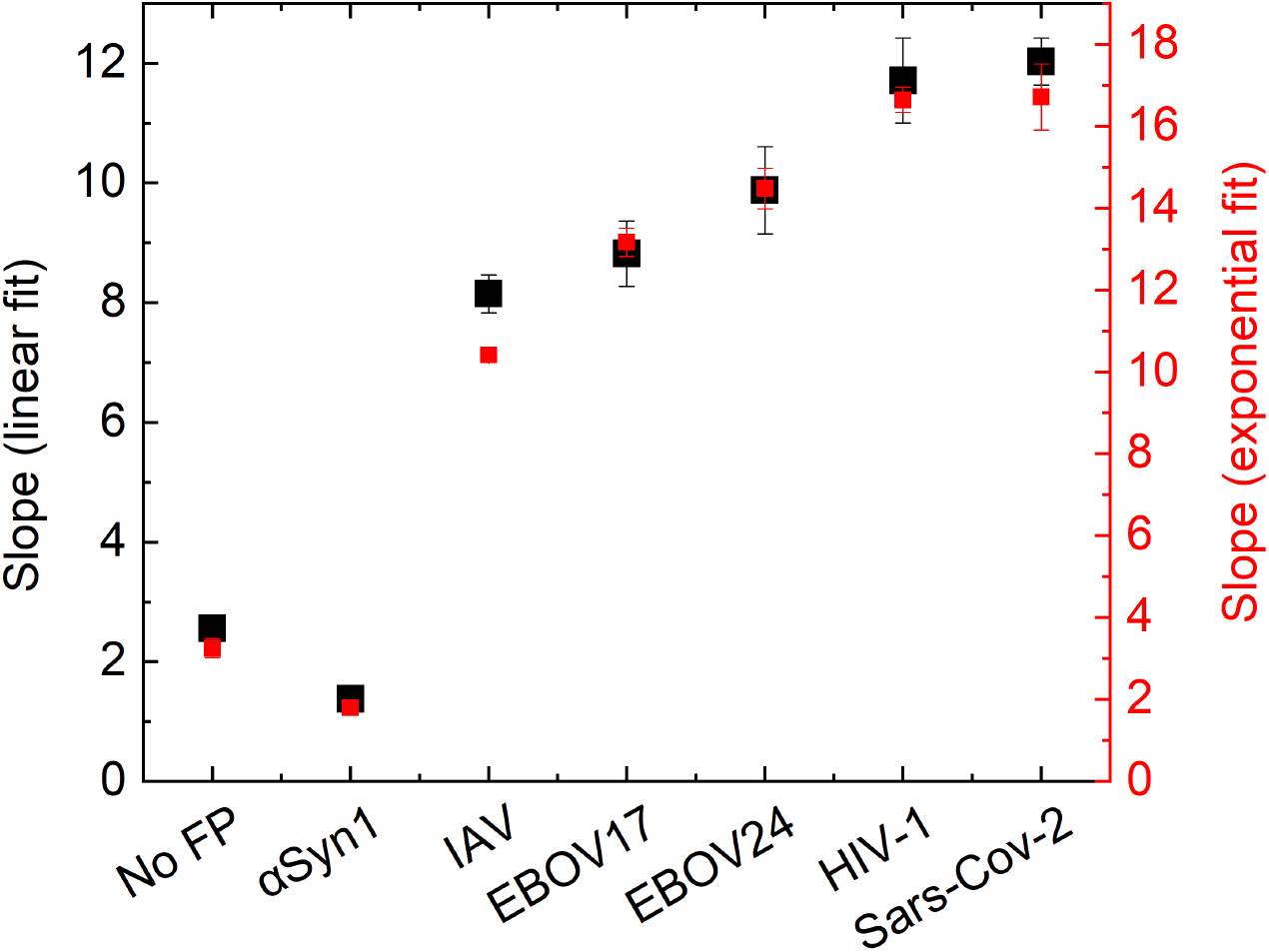
Comparison of fusion efficiencies for different fusion peptides and control samples. Shown are the slopes obtained from linear fits (black data) together with the initial slopes at *t* = 0 derived from exponential fits (red data) for each sample. Values correspond to the average of five independent experiments, and error bars represent the standard deviation of the mean.

Across both analysis methods, the studied VLPs showed comparable trends of fusion efficiency, whereas control systems lacking fusion peptides (No FP) or containing non-binding peptides (*α*Syn1) showed approximately 3–5 times lower efficiency than VLP-containing samples. Notably, the *α*Syn1 peptide consistently exhibited even lower fusion efficiency than the No FP condition. Among the viral systems, IAV VLPs showed the lowest fusion efficiency, while HIV-1 and SARS-CoV-2 displayed the highest values, corresponding to approximately 40% difference between the least and the most efficient system. Importantly, almost identical relative fusion efficiencies were detected between the exponential and linear fitting approaches even that the absolute values vary by about 25 %. This indicates that the extracted trends are robust with respect to the chosen analytical model.

These results strongly suggest that fusion peptides are not limited to passive membrane targeting but instead contribute to the fusion process itself. Their presence significantly enhances membrane fusion efficiency, most likely by promoting membrane insertion and inducing local perturbations that reduce the energetic barrier for lipid mixing. This interpretation is consistent with previous structural and biophysical studies proposing that fusion peptides destabilize target membranes during early fusion intermediates (*23*).

To further understand the observed differences between the peptide systems, the results were interpreted in the context of sequence-dependent physicochemical properties that influence peptide-membrane interactions. Variations in fusion efficiency can be linked to differences in the distribution of charged residues, charge patterning, and structural organization of the peptides (*8, 24*). In particular, the balance between hydrophobic segments and charged residues is known to critically determine membrane insertion probability (*25*). Fusion peptides containing extended, continuous hydrophobic stretches are more likely to exhibit a stronger membrane affinity to a neighboring membrane and a higher fusion activity, whereas interruptions in hydrophobic continuity or unfavorable charge clustering can impair membrane insertion.

The amino acid sequences analyzed in this study are summarized in Table 2, with charged hydrophilic residues highlighted in red. Comparative inspection reveals that viral fusion peptides maintain relatively continuous hydrophobic cores despite the presence of charged residues such as arginine (R), aspartate (D), glutamic acid (E), and lysine (K), which are mainly concentrated at one of the terminal regions. In contrast, the *α*Syn1 peptide exhibits a more dispersed distribution of charged residues, including at the terminal regions, which disrupts hydrophobic continuity and reduces membrane insertion efficiency. This structural feature is consistent with he observation that its negligible fusion activity is even lower than that of the control sample lacking fusion peptides.

Among the tested viral peptides, the IAV fusion peptide contains more frequent interruptions within its hydrophobic segment than other sequences, which might explain its comparatively reduced fusion efficiency. This observation is further supported by previous studies demonstrating that IAV fusion peptide activity is strongly pH-dependent, with acidic conditions enhancing membrane interaction by modulating charge states of the key residues (*26*). In addition, hydropathy analysis using the Kyte–Doolittle scale (*27*) were obtained and summarized in Table 2. Although GRAVY values near neutrality should be interpreted with caution, the distinctly negative hydropathy of *α*Syn1, combined with its disrupted hydrophobic organization, provides a reasonable explanation for its minimal fusion activity, which is even lower than the peptide-free control condition.

## Conclusion and Outlook

In this study, we investigated the functional role of viral fusion peptides in membrane fusion using a controlled microfluidic assay with model lipid bilayers. Although the anchoring function of fusion peptides is well established, their mechanistic contribution along the fusion pathway has remained unclear. Our results demonstrate that fusion peptides are not merely structural elements that position fusion proteins at the membrane interface but are active contributors to the fusion process itself.

A clear and consistent increase in the fusion efficiency was observed for enveloped VLPs compared to control systems lacking functional fusion peptides or containing non-binding peptides. These pronounced differences confirm that fusion peptides significantly enhance membrane fusion rather than acting as passive anchors. The observed trends further suggest that the fusion efficiency is strongly influenced by the ability of the peptides to interact with and perturb the target membrane.

Analysis of different viral systems supports a model in which sequence-dependent physicochemical properties, particularly hydrophobic segment length and charge distribution, govern membrane insertion and fusion activity. In this framework, HIV-1 and SARS-CoV-2 fusion peptides exhibit the most favorable characteristics for membrane insertion, while IAV fusion peptides show comparatively reduced efficiency, consistent with their interrupted hydrophobic profiles.

Future work should aim to resolve the molecular mechanisms of peptide-lipid interactions in more detail, including low pH, and peptide conformation collectively regulate fusion efficiency. In particular, combining microfluidic assays with structural and computational approaches could provide a more detailed mechanistic understanding of how fusion peptides modulate membrane remodeling during viral entry.

## Materials and Methods

### Material

The used lipids including 1,2-Dioleoyl-sn-glycero-3-phosphocholine (DOPC), 1,2-dioleoyl-sn-glycero-3-phosphoethanolamine (DOPE), 1,2-dioleoyl-sn-glycero-3-phospho-L-serine (DOPS), 1,2-distearoyl-sn-glycero-3-phosphate (sodium salt) (DSPA), 1,2-dioleoyl-snglycero-3-phosphoethanolamine-N-(lissamine rhodamine B sulfonyl) (ammonium salt) (18:1 Liss Rhod-PE), 1,2-dioleoyl-sn-glycero-3-phosphocholine-N-(Cyanine 5) (18:1 Cy5-PC), N-octadecanoyl-D-erythro-sphingosylphosphorylcholine, and cholesterol, were all purchased from Avanti Polar Lipids. 1,2-Dioleoyl-sn-glycero-3-phosphoethanolamine-N-(carboxyfluorescein) (ammonium salt) (FITC-PE) was obtained from Santa Cruz Biotechnology (ChemCruz). Dimethyl sulfoxide 99+% (DMSO) and potassium chloride (KCl) were purchased from Thermo Fisher and Sigma-Aldrich, respectively. The aqueous KCl buffer solution was prepared using ultrapure water (Thermo Fisher). All the studied peptides were synthesized by ProteoGenix S.A.S., France.

### Fabrication of the microfluidic device

For fusion experiments, a microfluidic device was fabricated in Sylgard 184 featuring a long straight channel with circular areas at both ends for inlets and outlet. The total dimensions of the channel are 1 mm in width, 100 µm in height and 10 mm in length. The microfluidic devices were fabricated using standard soft-lithographic protocols. For that, first masters were prepared in SU8-100 photoresist (Kayaku advanced material) on silicon wafers obtained from Si-Mat -Silicon Materials e.K.. The photoresist was spin-coated (B.L.E manufacturer, delta 10 model) on the Si-wafer until full coverage. Subsequently, the coated wafers were soft baked at 65 °C and 95 °C for about one hour to evaporate the solvent and increase the density of the film. After that, the photoresist-coated wafer was exposed through a transparency photomask (Micro Lithography Services) to UV light with a peak wavelength of about 400 nm and an intensity of 20 mW*/*cm^2^ for10 s. This step is followed by a 6 min post-exposure bake at 65 °C and 12 min at 95 °C to crosslink the exposed parts of the film. Finally, the wafers were immersed and gently shaken in the developer solution (Mr-dev600, microresist technology), which dissolves the non-crosslinked parts of the photoresist. All the procedure was carried out in a cleanroom environment to avoid undesired UV exposure and to minimize the risk of contamination.

The thus produced SU-8 structures serve as a master for the fabrication of the microfluidic device from PDMS. For that, the two components of the commercial elastomer kit “Sylgard 184” (Dow chemical) were mixed and degassed in a vacuum chamber. Subsequently, the degassed mixture was poured over the SU-8 master with a layer thickness of about 4 mm and fully cured at 85 °C for 3 h. The cured PDMS was carefully detached from SU-8 master and inlets and outlets were punched into the Sylgard 184 structure at both ends of the channel using a Harris Uni-core 1.0 mm puncher, and plasma bonded (Diener Electronics, type Femto) to a thin glass cover slide (thickness 0.13-0.16 mm). To strengthen the bonding of the Sylgard 184 to the glass, the device was placed on a hotplate at 85 °C for 2 h.

### Lipid Bilayer Formation and Characterization

Supported lipid bilayers were formed using LUVs, which were prepared through extrusion method using Avanti mini extruder. The composition of the supported lipid bilayer used in this study can be found in Table 1. It includes 2 mol% of fluorescent labelled PC (Cy5-PC) having a maximum excitation and emission wavelengths of 647 nm and 665 nm, to facilitate the observation of vesicles by fluorescence microscopy. To prepare the solution that is used to form the vesicles, initially each component was individually dissolved in chloroform, in their original quantity, and stored at −20 °C. To form the bilayer LUVs, specific amounts of each component were combined according to the molar ratios in Table 1 to reach a total concentration of 1 mg/mL. The chloroform was evaporated under vacuum for about one hour. The dry lipid film was dissolved in 150 mM KCl buffer at room temperature for 30 min and thoroughly mixed by pipetting and vortexing to achieve a uniform lipidic solution. This solution was passed 10 times through the mini extruder with a 0.1 µm polycarbonate membrane above the phase transition temperature, producing LUVs with an approximate diameter of 100 µm. Supported lipid bilayer on a glass substrate, i.e. the glass bottom of our straight microfluidic channel, were formed using the LUV solution that were prepared as described, and injecting them into the microfluidic device by a syringe pump. The initial flow rate was set at approximately 100 µL/min and after one minute, once the device was fully filled with the vesicle solution, the flow rate was reduced to 10 µL/min and maintained for about 10 min. After stopping the flow, the bilayer was incubated for 10 minutes. Following the incubation time, another injection of LUVs was carried out with a flow rate of 10 µL/min for 5 min to ensure a defect-free bilayer and complete coverage of the channel. Finally, the channel was thoroughly rinsed with 150 mM buffer solution at a flow rate of 100 µL/min for 2 min, before proceeding with the fusion experiments.

The microscopy observations were all performed by an inverted Axio Observer 7 (Carl Zeiss) fluorescence microscope and all steps for the preparation of the supported bilayer were performed at room temperature. Fluorescence Recovery after Photobleaching (FRAP) measurements were used to determine the lateral mobility in the formed supported lipid bilayers. For that, a defined region of interest was photobleached using high-intensity LED illumination through a 50 µm pinhole aperture for approximately 30 s. Fluorescence recovery was subsequently monitored at low illumination intensity to minimize additional photobleaching. Time-lapse images were recorded until recovery reached a plateau, as shown in Fig 2.

### Fusion experiments

For fusion experiments, the representative peptides from Orthomyxoviridae family: IAV (Influenza A virus); Coronaviridae family: SARS-CoV-2 (Severe acute respiratory syndrome coronavirus 2); Retroviridae family: HIV-1 (Human immunodeficiency virus type 1); and Filoviridae family: EBOV (Ebola Virus) were chosen. EBOV17 and EBOV24 correspond to fusion peptides from ebola virus of different lengths that were used in the experiments (*28–31*). As controls, a known human-sourced non-binder peptide *α*-Synuclein1 (Mut5) and LUVs without any fusion peptides were used (peptide sequences are listed in Table 2) (*32*). All of the peptides were synthesized by ProteoGenix S.A.S., France, with the purity of more than 90%. These peptides were dissolved in DMSO and stored at −80 °C according to supplier’s guideline. Virus like particles (VLPs) were prepared using the same protocol as for the LUVs that were used to form the supported bilayer. In contrast to the LUVs for bilayer production, however, a modified lipid composition is used that is closer to the virus cell membranes and contains 2 mol% of fluorescent labeled PE (Rhod-PE) (Table 1) (*22*). Rhod-PE has a maximum excitation and emission wavelengths of 560 nm and 583 nm, respectively, which differ sufficiently from the wavelengths of Cy5-PC, enabling the distinction of VLPs from the bilayer. Before a fusion experiment, small aliquots of the peptide solutions were added to the vesicle solution to achieve a 10:1 molar ratio of peptides to LUVs. DMSO was always kept below 1 vol% of the buffer to prevent any effect on the vesicles’ surface properties or fusion capability (*28, 33, 34*).

At the beginning of a fusion experiment, the VLP solution was injected with a syringe pump into the microfluidic channel containing the previously formed supported lipid bilayer. To achieve a temporal resolution of the fusion activity despite the strong background fluorescence from the VLPs, the process was conducted in multiple cycles. Each cycle included an initial injection of VLP at a flow rate of 10 µL/min for 20 s, followed by rinsing the channel with 150 mM KCl buffer at 100 µL/min for 2 min. After each rinsing step, a fluorescent image was captured by the fluorescence microscope. This injection and observation cycle was repeated several times to obtain the fusion efficiency of the different VLPs and control vesicles, as determined by the fluorescent intensity.

## CRediT authorship contribution statement

M. Mohammadian: Investigation, Formal analysis, Data curation, Software, Writing – original draft. R. Seemann: Conceptualization, Funding acquisition, Supervision, Writing– review and editing.

## Acknowledgement

We would like to thank Dr. Jean-Baptiste Fleury, who came up with the idea of this project. Financial support is acknowledged by the Deutsche Forschungsgemeinschaft (DFG), Germany, SFB1027 (project B4).

